# Liver Endothelium Microenvironment Promotes HER3-mediated Cell Growth in Pancreatic ductal adenocarcinoma

**DOI:** 10.1101/2022.06.24.497507

**Authors:** Wei Zhang, Michel’le Wright, Moeez Rathore, Mehrdad Zarei, Ali Vaziri-Gohar, Omid Hajihassani, Ata Abbas, Hao Feng, Jonathan Brody, Sanford D Markowitz, Jordan Winter, Rui Wang

## Abstract

∼90% metastatic pancreatic ductal adenocarcinoma (mPDAC) occurs in the liver, and the 5-year survival rate for patients with mPDAC is only at 3%. We previously reported that liver endothelial cells (ECs) secreted soluble factors to promote colorectal cancer cell survival in a paracrine fashion. However, the effects of liver ECs on mPDAC have not been elucidated. In this study, we used primary ECs from non-neoplastic liver tissues. We treated PDAC cells with conditioned medium (CM) from liver ECs, with CM from PDAC as controls, and determined that liver EC-secreted factors increased PDAC cell growth. Using an unbiased receptor tyrosine kinase array, we identified human epidermal growth factor receptor 3 (HER3, also known as ErbB3) as a key mediator in EC-induced growth in PDAC cells that have HER3 expression (HER3 +ve). We found that EC-secreted neuregulins activated the HER3-AKT signaling axis, and that depleting neuregulins from EC CM or blocking HER3 with an antibody, seribantumab, attenuated EC-induced proliferation in HER3 +ve PDAC cells, but not in cells without HER3 expression. Furthermore, we determined that EC CM increased PDAC xenograft growth *in vivo*, and that seribantumab blocked EC-induced growth in xenografts with HER3 expression. These findings elucidated a paracrine role of liver ECs in promoting PDAC cell growth, and identified the HER3-AKT axis as a key mediator in EC-induced functions in PDAC cells with HER3 expression.

**Implications:** over 70% mPDAC express HER3. This study suggests the potential of using HER3-targeted therapies for treating patients with HER3 +ve mPDAC.

## Introduction

Pancreatic ductal adenocarcinoma (PDAC) is the third-leading cause of cancer-related death in the United States and is predicted to become the second over the next decade (1). Five-year survival is just 39% for ‘curable’ PDAC after surgery, on par with metastatic disease for other tumor types. Over 50% of patients with PDAC develop distant metastases [mPDAC, ∼90% found in the liver (2)] and only have a five-year survival rate at 3% (3). All patients are recommended to receive systemic chemotherapy, regardless of stage, yet these treatments are only marginally effective (4-7). Therefore, novel therapeutic strategies are urgently needed to treat patients with this devastating disease, especially with mPDAC.

Extensive preclinical studies focused on the intra-tumor stromal cells such as fibroblasts in PDAC but have not made an impact in the clinic (8-12). On the other hand, one key component of the tumor microenvironment that has been overlooked is the surrounding host organ. The liver is the most common site of distant metastases in mPDAC. It has a unique endothelium-rich microenvironment, with up to ∼40% of liver stroma composed of endothelial cells (ECs) (13-15). Liver ECs are known to interact with adjacent cells and play a key role in liver physiology and pathology. Previous studies reported that interactions between circulating tumor cells and liver ECs promote colonization of cancer cells in the liver, and that ECs induce immune tolerance and surveillance escape in the liver (16). Meanwhile, studies from our laboratory reported that liver ECs secrete soluble factors to promote colorectal cancer (CRC) cell survival in a paracrine fashion (17-20), partially by activating human epidermal growth factor receptor 3 (HER3, also known as erb-b2 receptor tyrosine kinase 3, ErbB3). HER3 is a member of the HER receptor tyrosine kinase (RTK) family, which consists of EGFR, HER2, HER3 and HER4. Different from other HER receptors, HER3 only has a pseudokinase domain with weak activity and predominantly relies on dimerizing with other HER family receptors for activation. When HER3 binds to its only known ligand, neuregulins (NRGs, also known as heregulins), dimerization of HER3 and other HER receptors (most commonly with HER2) follows, which then causes trans-phosphorylation and activation of HER3 (21). In CRC and other types of cancer, HER3 activation leads to the activation of downstream pathways including AKT and MAPK that have significant pro-survival roles.

In PDAC, TCGA data indicate that HER3 mRNA levels are higher in tumors compared to normal tissues (22). Interestingly, immunohistochemical (IHC) staining revealed that HER3 proteins are expressed in less than 30% of primary tumors and in nearly 70% of PDAC liver metastases, and high protein levels of HER3 correlate with poor overall survival in patients with PDAC (23,24). Moreover, limited preclinical studies showed that the canonical NRG-induced HER2-HER3 activation promoted PDAC cell growth, and that inhibiting HER2 or HER3 with preclinical compounds decreased PDAC cell growth (25,26). However, the effects of liver EC microenvironment on PDAC cell functions and the role of HER3 in mediating EC-PDAC crosstalk has not been determined. In this study, we aim to determine the role of liver ECs in mediating PDAC cell functions, and identify the involved signaling pathway(s). We demonstrated that conditioned medium (CM) containing EC-secreted factors significantly increased PDAC cell growth, and found that EC-secreted NRG activated PDAC-associated HER3 and AKT for promoting cell growth in PDAC cells with HER3 expression (HER3 +ve).

Blocking HER3, either by NRG depletion from EC CM or by the HER3-specific antibody, seribantumab, attenuated EC-induced HER3-AKT activation and cell growth in HER3 +ve PDAC cells, whereas seribantumab had no effects on PDAC cells without HER3 expression (HER3 -ve). Furthermore, we used a proof-of-principle subcutaneous (subQ) xenograft model to determine that EC CM promoted PDAC xenograft growth, and seribantumab blocked EC-induced HER3-AKT activation and growth in xenografts. Overall, our results demonstrate a paracrine role of liver ECs in promoting PDAC cell growth, and identify the HER3-AKT signaling axis as a key mediator of EC-induced functions in HER3 +ve PDAC cells. As seribantumab had significant on-target inhibition effects in preclinical and clinical studies in other types of cancer (18,27-31), our findings highlight a potential strategy of using HER3 antibodies for treating patients with PDAC, with HER3 expression being a potential predictive marker for HER3-targeted therapies.

## Materials and Methods Cell culture

The established human PDAC cell lines BxPC-3, Capan-2, PANC-1 and MIA-PaCa2 were purchased from ATCC (Manassas, VA, USA). Human primary liver ECs (EC1 and EC9) lines were isolated and established in our laboratory using MACS microbead-conjugated anti-CD31 antibodies and separation columns (Miltenyi Biotec, Bergisch Gladbach, Germany), as described previously (18-20,32,33). All PDAC cells were cultured in DMEM (Sigma-aldrich, St. Louis, MO, USA) supplemented with 5% FBS (Atlanta Biologicals, Atlanta, GA), vitamins (1x), nonessential amino acids (1x), penicillin-streptomycin antibiotics (1x), sodium pyruvate (1x), and L-glutamine (1x), all from Thermo Fisher Scientific/Gibco (Grand Island, NY, USA). Primary human liver ECs were cultured in Endothelial Cell Growth Medium MV2 (PromoCell, Heidelberg, Germany) supplemented with 10% human serum (Atlanta Biologicals) and antibiotic-antimycotic (1x, Thermo Fisher Scientific/Gibco). Primary ECs were used within 10 passages (approximately 1 week per passage). All other established PDAC cell lines were used within ∼20 passages after receipt.

Cell line mutations were determined in previous studies (20), or from public databases including the Cancer Cell Line Encyclopedia (CCLE) and the Catalogue of Somatic Mutations in Cancer (COSMIC). Mycoplasma contamination and short tandem repeat (STR) authentication tests for all cell lines are done every 6 months (last tested on 9/8/2021 by the Genomics Core at Case Comprehensive Cancer Center, Case Western Reserve University). For primary liver EC lines, genomic DNA samples from original tissues were used for the STR authentication.

### Conditioned medium (CM)

3 × 10^5^ of PDAC cells or ECs were seeded in T25 culture flasks for two days. After, the cells were washed two times with 1X PBS and then cultured in 3 mL DMEM with 1% FBS (1×10^5^ cells/mL) for 48 hours. CM were harvested and centrifuged at 4,000 g for 10 minutes to remove cell debris. CM from each PDAC cell line were used as controls. For CM treatment, all PDAC cells were incubated in DMEM medium with 1% FBS overnight, and then treated with control or EC CM for the indicated time.

### Reagents

The fully humanized IgG2 anti-HER3 antibody seribantumab (previously known as MM-121) was provided by Elevation Oncology Inc. (New York, NY, USA). HER2 antibody trastuzumab is from MedChemExpress LLC. (Cat#: HYP-9907, Monmouth Junction, NJ). The control human IgG antibody for *in vitro* and *in vivo* studies was from Invitrogen (Carlsbad, CA, USA). 250 μg/mL seribantumab and 50 μg/ mL trastuzumab were used for all *in vitro* studies. Recombinant human NRG (rhNRG) was from BioVision (Cat# 4711-10, Waltham, MA) and used at the indicated levels. Human *NRG1*-specific siRNAs (5’-UCGGCUGCAGGUUCCAAAC, and 5’-GGCCAGCUUCUACAAGCAU) and a validated scrambled control siRNA were obtained from Sigma-Aldrich (St. Louis, MO).

### Receptor tyrosine kinase (RTK) array

The RTK array kit was purchased from R&D Systems (Cat# ARY001B, Minneapolis, NE) and the assay was performed according to the manufacturer’s instructions. In brief, 0.5 ×10^6^ PDAC cells were incubated in DMEM medium with 1% FBS overnight, and then treated with control or EC1 CM for 30 minutes. Cell lysates were prepared in lysis buffer from the kit and 300 μg total proteins from each group were loaded onto the membranes.

### Western blotting

PDAC cells were treated with CM (without or with antibodies) for 30 minutes. Cell lysates were processed and run through SDS-PAGE gel electrophoresis as described previously (18,34). An HRP-conjugated β-actin antibody was obtained from Santa Cruz Biotechnology and was used in 1:8000 dilution (Santa Cruz Biotechnology Cat# sc-47778 HRP, Santa Cruz, CA, USA). All other antibodies were obtained from Cell Signaling Technology and were used in a 1:1000 dilution (Beverly, MA, USA). Phospho-AKT S473 (Cell Signaling Technology Cat# 9271), total AKT (Cell Signaling Technology Cat# 9272), Phospho-HER3 Y1289 (Cell Signaling Technology Cat# 2842), total HER3 (Cell Signaling Technology Cat# 12708). Phospho-HER2 Y1248 (R&D Systems, Cat# AF1768-SP) and total HER2 (Neu C-3) (Santa Cruz, Cat# sc-377344). For each experiment, protein lysates were loaded into multiple gels with 50 µg total proteins per lane and processed simultaneously to separately probe for antibodies specific to phosphorylated proteins and total proteins. All membranes were probed with β-actin as loading controls and a representative loading control was used in the figures. Western blotting images were developed by enhanced chemiluminescent (ECL) substrate and autoradiography films (both from Thermo Fisher Scientific), and figures presented representative results from one experiment of at least three independent experiments.

### MTT assay

PDAC cells were seeded at 2,000 cells/well in 96-well plates, cultured in DMEM medium with 1% FBS overnight and then incubated in CM for 72 hours. Cells were pretreated with seribantumab (250 μg/mL), or/and trastuzumab (50 μg/mL) overnight, and then cultured in CM with seribantumab for 72 hours. Cell proliferation was assessed by adding MTT substrate (0.25% in PBS, Sigma-Aldrich) in DMEM medium with 1% FBS (1:5 dilution) for 1 hour at 37 °C. Cells were washed with 1x PBS and then added with 50 μL DMSO. The optical density of converted substrate was measured at 570 nm, and the relative MTT was presented as percent of control groups with cells treated with PDAC CM.

### PicoGreen assay

PDAC cells were seeded at 2,000 cells/well in 96-well plates, cultured in DMEM medium with 1% FBS overnight and then incubated in CM for 72 hours. After treatment, the culture medium was discarded and 100 μL molecular grade water was added to the cells for a 1-hour incubation and then added with 100 μL 1X PicoGreen (Invitrogen, Cat# P11496) in molecular grade water for another 1 hour. Fluorescence was measured at 480 nM by a microplate reader and presented as percent of control groups with cells treated with PDAC CM.

### Colony formation

PDAC cells were seeded in 6-well plates at a density of 8000 cells/well. Next day, cells were switched to DMEM medium with 1% FBS overnight, and then incubated in CM from PDAC or ECs for 72 hours. Then, cells were gently washed with cold 1x PBS and stained with 300 μL/well crystal violet (0.5% w/v crystal violet in 20% methanol) at room temperature for 30 minutes, followed by washing with water, air dry overnight, and imaged with Bio-Rad imager (ChemiDoc XRS+ System). Finally, crystal violet was eluted by 1% SDS solution in 300 μL /well DMSO. Absorbance was measured at 600 nM by microplate reader, and presented as density relative to control groups with PDAC CM.

### siRNA transfection

For each transfection, 2 × 10^5^ ECs were transiently transfected with 100 pmol siRNAs (control scrambled or *NRG*1-specific) plus 5 μL of lipofectamine in 150 μL OptiMEM medium overnight. Cells were recovered in DMEM medium with 5% FBS for 24-48 hours, and then subjected to generate CM, as described above. Two *NRG*1-specific siRNAs were mixed 1:1 for transfection.

### Xenograft tumor models

3 × 10^6^ BxPC-3 cells were injected subcutaneously (subQ) into the right flanks of athymic nude mice in 100 μL growth-factor-reduced Matrigel. After tumor burden was confirmed by a caliper, mice were randomized to four groups with equal average tumor sizes, and then treated with 3x concentrated PDAC or EC CM by subQ injection adjacent to implanted tumors once a week. IgG control antibody or seribantumab (5 mg/mouse) were delivered by intraperitoneal (IP) injection twice a week. Tumor volumes were measured by a caliper.

### Immunohistochemistry (IHC)

Xenografts were fixed in 4% paraformaldehyde, dehydrated in ethanol, and embedded in paraffin. Sections were cut 10 μm in thickness. Immunohistochemistry was done with antibodies against P-AKT (1:200; Cell Signaling Technology, Cat# 9271), P-HER3 (1:100; Cell Signaling Technology, Cat# 2842), human Ki67 (1:200, Cell Signaling Technology, Cat# 9449T). Isotype-matched immunoglobulin was used as a control. The HRP-conjugated anti-rabbit secondary antibody (Cell Signaling Technology, Cat# 8114p) were used with the DAB substrate kit (Vector Laboratories, Newark, CA, SK-4800) for staining. Stained sections were examined under a microscope (Keyence BZ-X810). The intensity of the staining was quantified by the QuPath software (NIH), and presented as density relative to control groups with PDAC CM.

### Statistical analysis

For *in vitro* assays, all quantitative data were reproduced in at least three independent experiments and multiple replications in each experiment. Groups were compared by a two-tailed student’s *t*-test and data were expressed as means -/+ standard error of the mean (SEM) with significance of P<0.01. For *in vivo* studies, one-way ANOVA was used for tumor volume changes over time, tumor sizes, and the quantification of P-HER3, P-AKT and Ki-67 staining between groups after tumor harvest. Data were presented as means -/+ standard deviation (SD) with a significance of P<0.05.

## Results

### CM from liver ECs increased PDAC cell growth

To determine the effects of liver ECs on PDAC cell growth, we used human primary liver ECs (EC1 and EC9) that we established from freshly resected non-malignant liver tissues (18,32). CM containing EC-secreted factors were harvested and added to human PDAC cells (BxPC-3, Capan-2, MIA-PaCa2 and PANC-1), with CM from PDAC cells themselves serving as control CM. First, the MTT assay was used to determine the effects of CM on cell viability. We found that compared to PDAC CM, CM from different liver EC lines significantly increased PDAC cell viability, with an over 2-fold increase in all four PDAC cell lines after 72 hours incubation (Fig 1A). To validate our findings, we then performed the PicoGreen assay, which detect double-strand DNA content in the cells, as a surrogate readout of the total number of cells. Similar to the results we observed from the MTT assay, CM from liver ECs significantly increased the number of viable cells compared to PDAC CM (Fig. 1B). Third, we performed the colony formation assay to further validate our findings. Results showed that, compared to PDAC CM, liver EC CM increased the number of colonies of cells by 2-2.5 fold in all four PDAC cell lines used (Fig. 1C, D). Together, data from three independent growth assays with multiple cell lines suggested that CM from liver ECs increased cell proliferation in PDAC cells.

**Figure 1.**
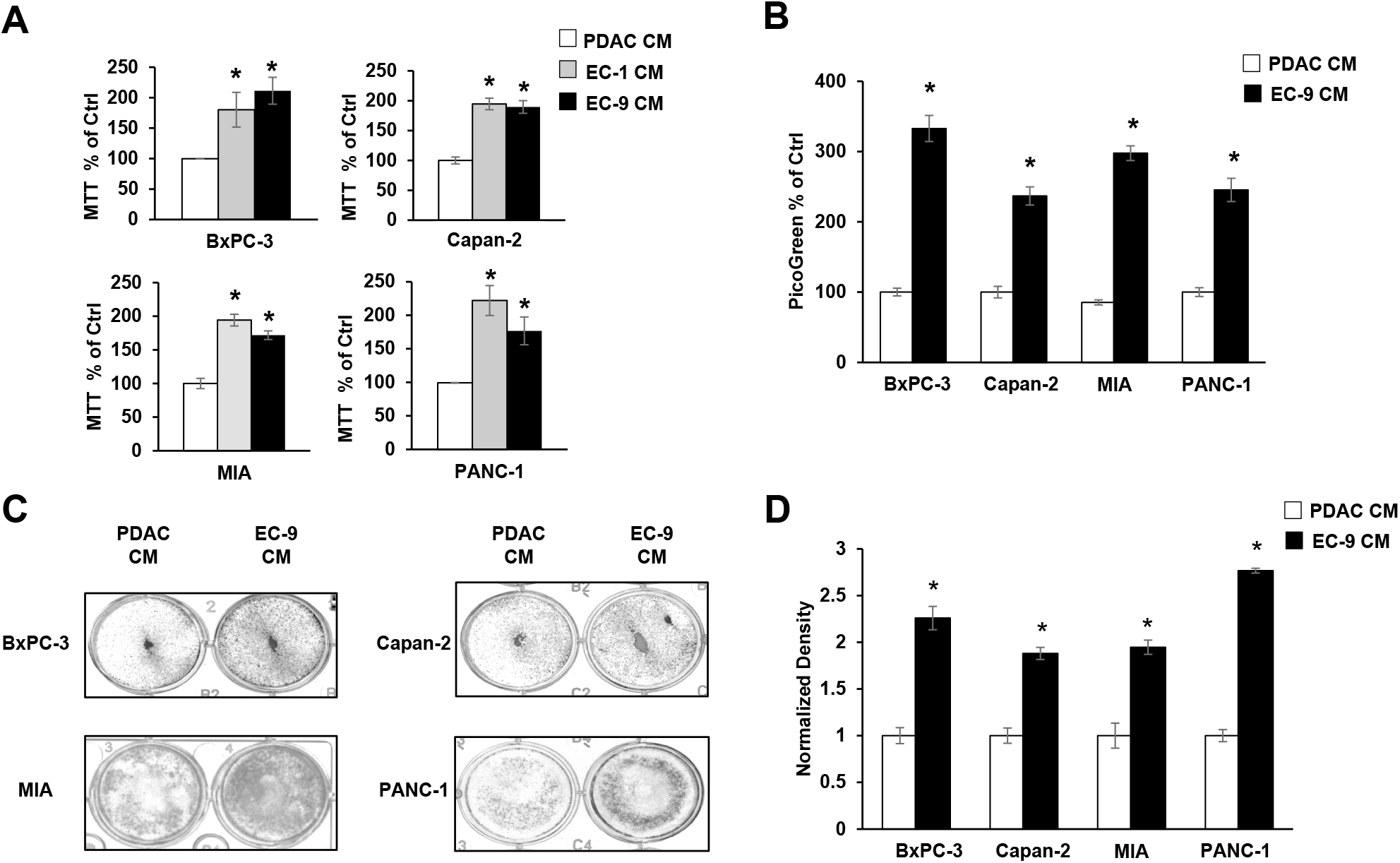
CM from Liver ECs increased PDAC cell growth *in vitro*. PDAC cells were incubated with control PDAC CM or CM from primary liver ECs (EC-1 or EC-9) for 72 hours. MIA, MIA-PaCa2 cells. **(A, B)** The MTT and PicoGreen assays, respectively, show that CM from liver ECs increased the cell viability and growth in four PDAC cell lines. **(C)** Representative images of PDAC cell colony formation after 72 hours in PDAC or EC-9 CM. **(D)** Quantification of the colony formation shows that EC CM increased the total amount of colonies. Mean +/-SEM of at least three experiments, *p<0.01 *t*-test compared to control groups with PDAC CM.

### CM from Liver ECs activated the HER3-AKT pathway in PDAC cells with HER3 expression

To understand the mechanism by which EC CM promoted PDAC cell growth, an unbiased receptor tyrosine kinase (RTK) array was used to assess the specific RTKs that were potentially activated by EC-secreted factors and were involved in promoting PDAC cell growth. We used BxPC-3 and PANC-1 cells and found that interestingly, these two PDAC cell lines had distinct RTK activation pattens when treated by EC CM for 30 minutes (Fig. 2A). In BxPC-3 cells, phosphorylation levels of HER2 and HER3 were the only two readouts that were noticeably increased by EC CM. In contrast, PANC-1 cells had no detectable HER2 and HER3 phosphorylation regardless of CM treatment, instead, the cells had increased levels of phosphorylation in insulin receptors (Insulin R). We then performed Western blotting to validate the findings from the RTK array. In addition to BxPC-3 and PANC-1 cells, we also used Capan-2 and MIA-PaCa2, and treated the PDAC cells with either EC CM or PDAC CM for 30 minutes (Fig. 2B). In agreement with the RTK array, we found that in both BxPC-3 and Capan-2 cells, EC CM increased the levels of phosphorylation in HER2 and HER3. Activation of AKT, an established downstream target of HER3 (21,35), was also detected. Western blotting also confirmed that in PANC-1 and MIA-PaCa2 cells, there was no detectable phosphorylation in HER2 and HER3 regardless of CM treatment. Moreover, we found that PANC-1 and MIA-PaCa2 cells had no detectable HER3 expression, even though HER2 expression was detected in these cells. However, regardless of HER3 expression, EC CM activated AKT in PANC-1 and MIA-PaCa2 cells. We attempted to validate the EC-induced phosphorylation of insulin receptors in PANC-1 cells observed from the RTK array. However, we could not detect consistent activation in insulin receptors after multiple repeats in PANC-1 and MIA-PaCa2 cells, suggesting insulin receptors may not be activated by EC CM. Of note, the levels of total and phosphorylated HER4, the other member of the HER receptor family, were not detected in the present study, or by others in previous studies (36). PANC-1 and MIA-PaCa2 cells reportedly have low levels of HER3 expression (25,37), and patient PDAC tissues have differential protein levels of HER3 (24). In agreement with those findings, our data confirmed that PDAC cells have heterogeneous expression profiles of HER3. Furthermore, our data suggest that the liver EC microenvironment activates the HER2-HER3 signaling axis in PDAC cells with HER3 expression (HER3 +ve cells) to induce the downstream pro-survival pathway AKT. The mechanism(s) by which liver ECs activate AKT in PDAC cells without HER3 expression (HER3 -ve cells, such as PANC-1 and MIA-PaCa2) remain unknown, and are being investigated in a separate study in our laboratory.

**Figure 2.**
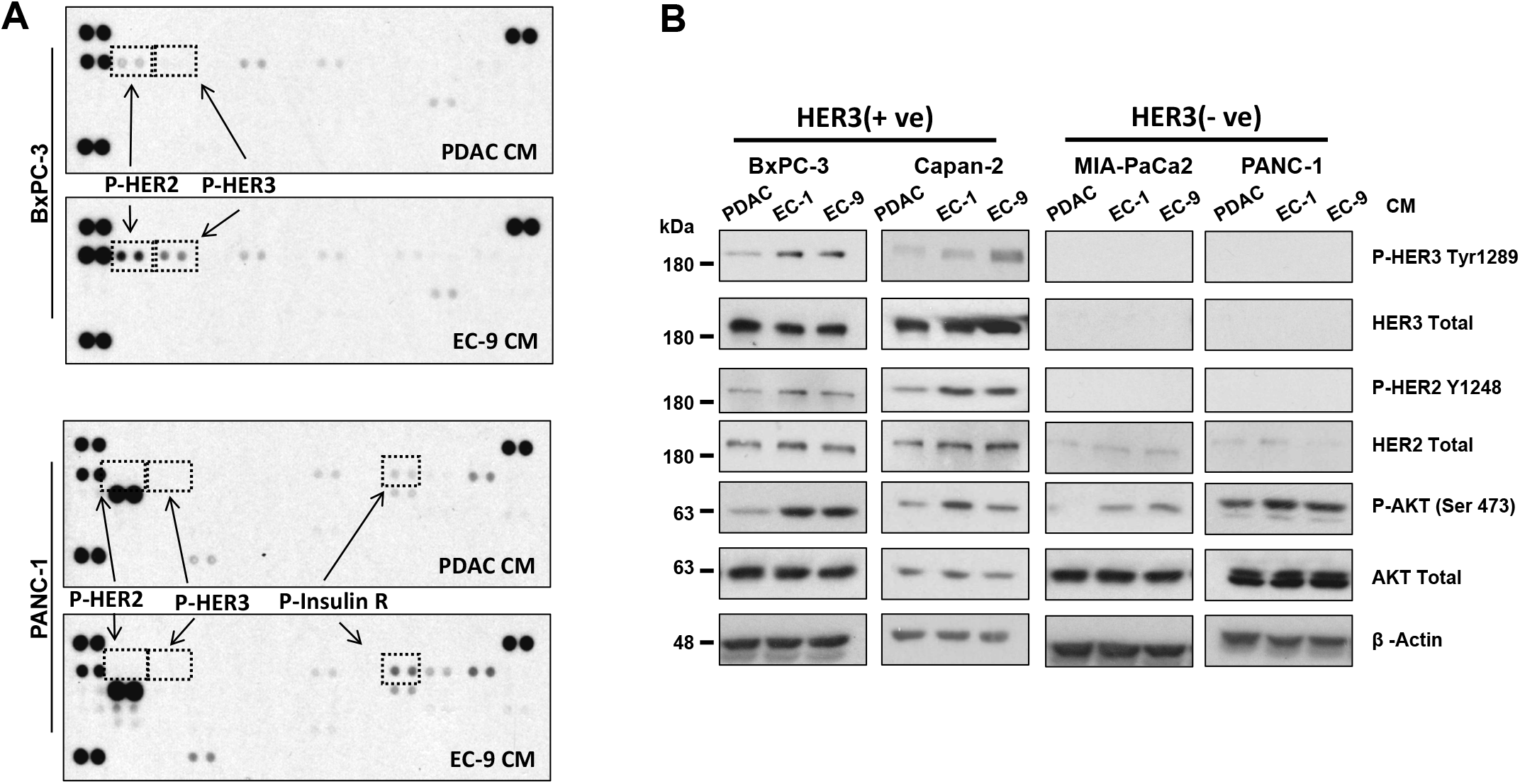
CM from liver ECs increased HER2-HER3 phosphorylation in PDAC cells with HER3 expression. PDAC cells were treated with CM for 30 minutes. **(A)** The RTK array shows phosphorylation of RTKs in BxPC-3 (upper) or PANC-1 (lower) cells treated by PDAC CM or EC-9 CM, with marked phosphorylation for HER2 (P-HER2), HER3 (P-HER3), and insulin receptor (P-Insulin R). **(B)** Western blotting shows that CM from ECs (EC-1 or EC-9) increased AKT phosphorylation in all cell lines, but only increased HER2-HER3 phosphorylation in BxPC-3 and Capan-2 cells. Total levels of HER3, HER2, AKT, and β-actin were used as loading controls. Data represent the results of at least three independent experiments.

### Liver EC-secreted NRGs activated HER3-AKT and promoted PDAC cell growth

To date, the only identified ligand for HER3 activation is NRG (21), which is known to be secreted by ECs (38). As EC CM activated HER3 in PDAC cells, we sought to determine whether the EC-induced HER3 activation and increased growth in PDAC cells were mediated by NRGs. First, we treated PDAC cells with recombinant human NRG (rhNRG) in PDAC CM, with EC CM treatment in parallel for comparison. Data from Western blotting (Fig. 3A) showed that EC CM activated HER3-AKT in BxPC-3 and Capan-2 cells (HER3 +ve), as expected, and rhNRG also induced HER3-AKT activation in these cells. Coupled with that, data from the MTT assay (Fig. 3B) showed that rhNRG promoted PDAC cell growth in a dose-dependent manner in these two cell lines. In contrast, rhNRG did not activate AKT in PANC-1 and MIA-PaCa2 cells (HER3 -ve), and had no effect on cell growth (in MIA-Paca2 cells, Suppl. Fig. 1), confirming that these two cell lines do not have a functional HER3 signaling axis. On the other hand, EC CM activated AKT and increased proliferation in these cells, further supporting the notion that ECs promoted PDAC cell growth in these cells via HER3-independent mechanism(s).

**Figure 3.**
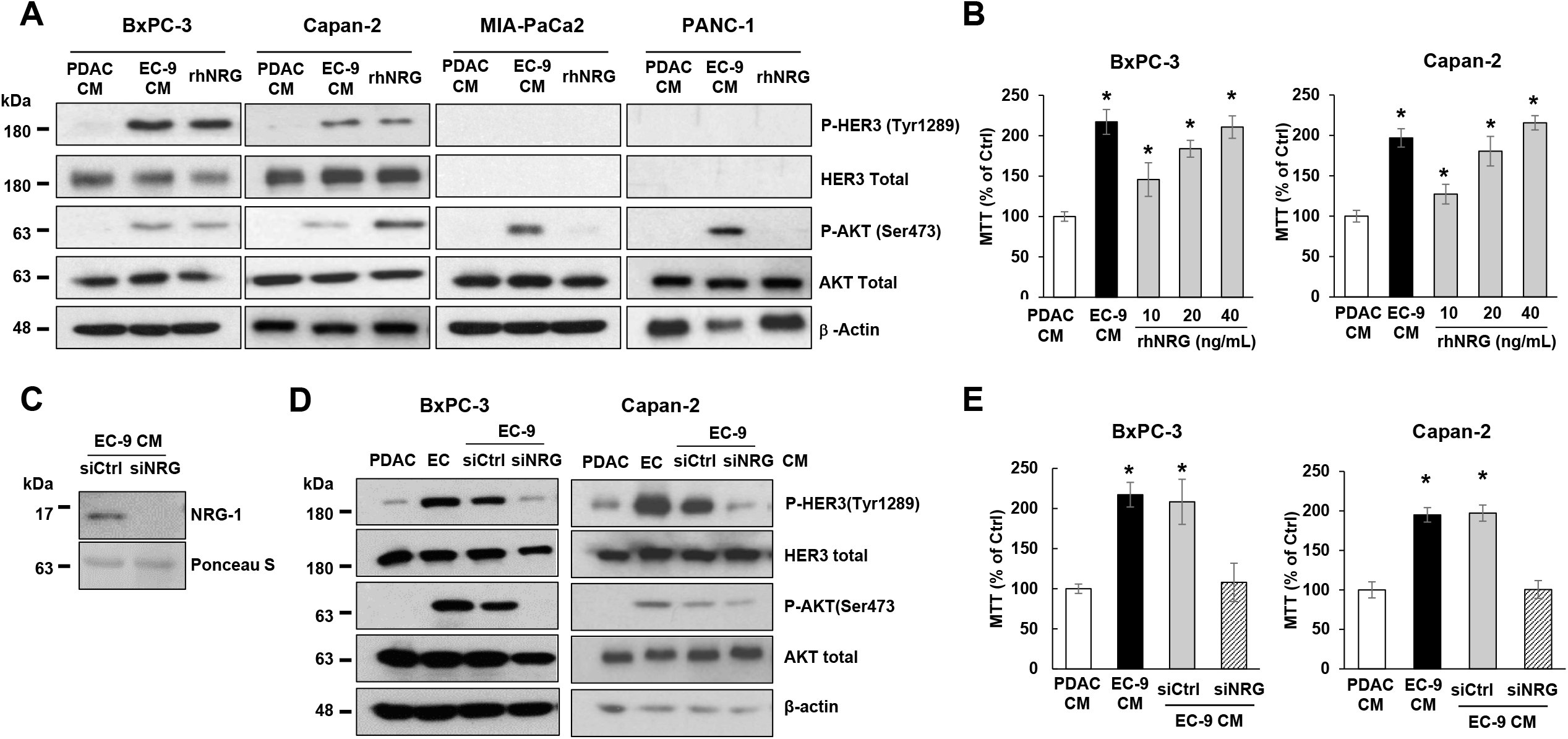
EC-secreted NRG activated HER3-AKT and promoted cell growth in PDAC cells with HER3 expression. PDAC cells were incubated in control PDAC CM, EC-9 CM, or recombinant human NRG (rhNRG) in PDAC CM. **(A)** Western blotting shows that HER3 and AKT were activated by EC CM and rhNRG in BxPC-3 and Capan-2 cells (HER3 +ve), and that rhNRG did not activate AKT in MIA-PaCa2 and PANC-1 cells (HER3 - ve). **(B)** The MTT assay shows that adding different doses of rhNRG in PDAC CM increased the cell viability in BxPC-3 and Capan-2 cells (HER3 +ve), with EC-9 CM as positive control and PDAC CM as negative controls. **(C)** Western blotting shows that transfecting ECs with NRG-specific siRNA (siNRG) depleted NRGs from EC CM, and that **(D)** NRG-depleted EC CM did not activate HER3-AKT in BxPC-3 and Capan-2 cells. **(E)** The MTT assay shows that NRG-depleted EC CM (by NRG-specific siRNAs) attenuated EC-induced PDAC cell viability. For Western blotting data, total levels of HER3, AKT, and β-actin were used as loading controls. Data represent the results of at least three independent experiments. For MTT assays, mean +/- SEM of at least three experiments, *p<0.01 *t*-test compared to control groups with PDAC CM.

To further determine that NRG is the key EC-secreted factor for activating HER3 and promoting growth in PDAC cells, we depleted NRGs from EC CM by knocking down NRG expression in liver ECs with *NRG1*-specific siRNAs. First, we confirmed that liver ECs expressed and secreted NRGs, and that *NRG1*-specific siRNAs decreased the levels of secreted NRGs in CM from liver ECs (Fig. 3C). Then, we applied NRG-depleted EC CM to HER3 +ve PDAC cells, with un-modified PDAC and EC CM as controls. We found that while un-modified EC CM containing NRGs activated HER3-AKT and promoted growth, as expected, NRG-depleted EC CM had no effect on HER3-AKT or cell growth in PDAC cells (Fig. 3D, E). We further validated our findings by depleting NRGs from EC CM with NRG-specific antibodies, and found that immuno-depleting NRG from EC CM attenuated the EC-induced HER3-AKT activation in PDAC cells (Suppl. Fig. 2). In summary, EC-secreted NRG is the key soluble factor for activating HER3-AKT and promoting cell growth in PDAC cells.

### HER3 inhibition blocked EC-induced cell growth in HER3 expressing PDAC cells

To determine the role of HER3 in mediating EC CM-induced AKT activation and PDAC cell growth, we blocked HER3 in PDAC cells with seribantumab, a humanized HER3-specific antibody that demonstrated significant HER3 inhibition in preclinical and clinical studies in other types of cancer (32,39,40). We first determined that seribantumab, at a level used in previous *in vitro* studies (250 μg/ml) (18,32,39), significantly blocked EC-induced HER3-AKT activation and subsequent cell growth in HER3 +v cells (BxPC-3 and Capan-2 cell, Fig. 4A, B). In contrast, seribantumab had no effect on EC-induced AKT activation or cell growth in HER3 -ve cells (PANC-1 and MIA-PaCA2, Fig. 4C, D). These findings suggest that in HER3 +ve PDAC cells, HER3-AKT is the key mediator of EC-induced PDAC cell growth. Moreover, seribantumab only blocked EC-induced growth in HER3 +ve cells, confirming that this antibody is specific for its target and suggesting that HER3 expression can be a potential predictive marker (or recruiting criteria) for treating patients with PDAC with seribantumab.

**Figure 4.**
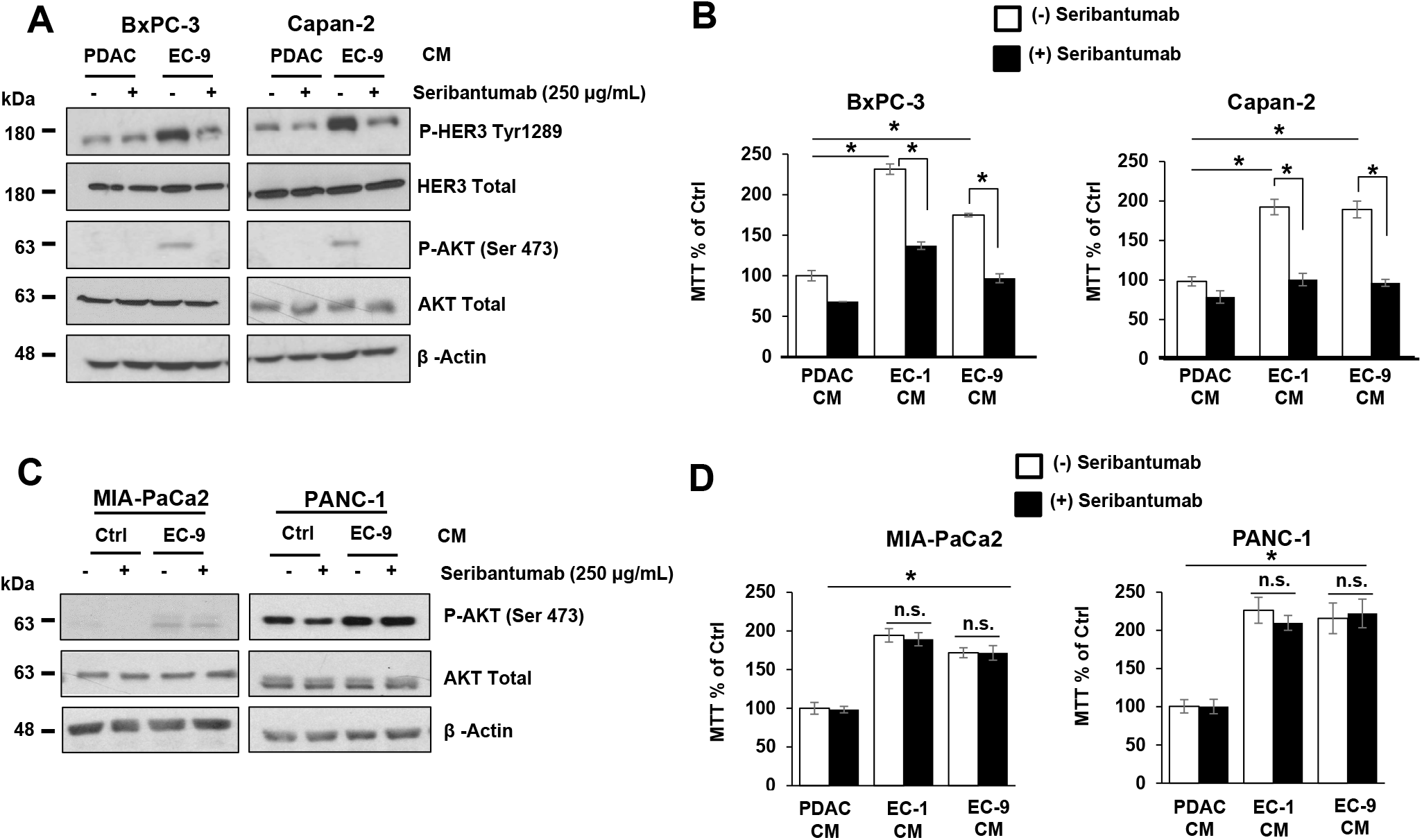
HER3 antibody seribantumab blocked liver EC-induced HER3-AKT activation and cell growth in PDAC cells with HER3 expression. PDAC cells were incubated in control PDAC CM or CM from different primary liver ECs (EC-1 or EC-9) either in the presence or the absence of the HER3 antibody seribantumab (250 µg/mL). **(A, B)** The Western blotting and MTT assay, respectively, show that seribantumab blocked liver EC CM-induced HER3-AKT phosphorylation and cell viability in BxPC-3 and Capan-2 cells (HER3 +ve). **(C, D)** The Western blotting and MTT assay, respectively, show that seribantumab did not block liver EC CM-induced AKT phosphorylation or cell viability in MIA-PaCa2 and PANC-1 cells (HER3 -ve). For Western blotting data, total levels of HER3, AKT, and β-actin were used as loading controls. Data represent the results of at least three independent experiments. For MTT assays, mean +/- SEM of at least three experiments, *p<0.01 *t*-test compared to control groups with PDAC CM without seribantumab.

Our data suggest that EC-secreted NRGs activated the canonical HER2-HER3 signaling pathway in PDAC cells. Since prior preclinical studies used another preclinical HER3-specific antibody and showed that the combination of HER2- and HER3-targeted therapies was superior to monotherapies in PDAC cells (25), we assessed the effects of combining seribantumab with trastuzumab, a HER2-specific antibody widely used in the clinic, on EC-induced PDAC cell functions. We first validated that trastuzumab was able to block EC-induced HER2, HER3, and AKT activations in BxPC-3 cells (Suppl. Fig. 3). Then we treated BxPC-3 and Capan-2 cells (HER3 +ve) with seribantumab or/and trastuzumab and assessed cell growth by the MTT assay (Fig. 5). Seribantumab blocked EC-induced cell growth, as expected, and trastuzumab blocked EC-induced PDAC cell growth to a lesser extent. Interestingly, we found that the combination of seribantumab and trastuzumab led to similar levels of growth inhibition compared to seribantumab alone, suggesting that HER2 inhibition did not augment the anti-growth effects of seribantumab. As a result, the effect of HER2 inhibition, either alone or in combination with seribantumab, was not assessed in subsequent *in vivo* studies.

**Figure 5.**
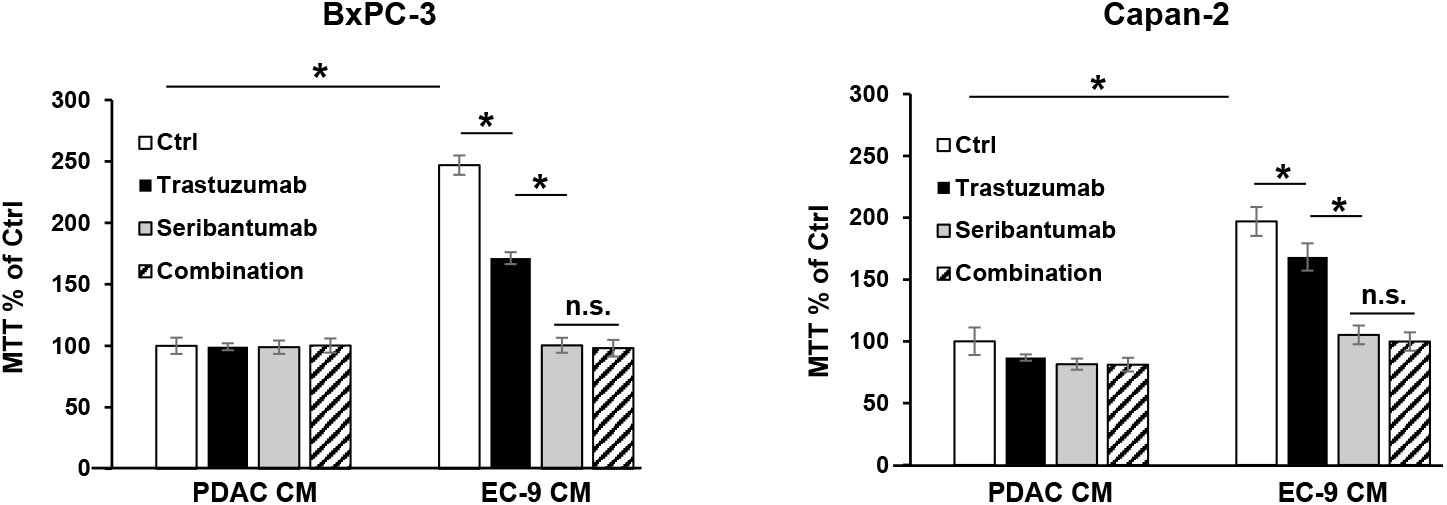
HER2 antibody trastuzumab did not augment the effects of seribantumab on blocking PDAC cell growth. BxPC-3 and Capan-2 cells (HER3 +ve) were incubated in control PDAC CM or CM from primary liver ECs (EC-9) either in the presence or the absence of the HER2 antibody trastuzumab (50 µg/mL) or the HER3 antibody seribantumab (250 µg/mL). The MTT assays showed that trastuzumab modestly blocked EC-induced PDAC cell viability, and that seribantumab completely blocked EC-induced PDAC cell viability. The combination of trastuzumab and seribantumab had similar inhibition effect on PDAC cell viability compared to seribantumab alone. Mean +/- SEM of at least three experiments, *p<0.01 *t*-test compared to control groups with PDAC CM without seribantumab, between adjacent groups. n.s., not statistically significant.

### HER3 inhibition blocked liver EC-induced PDAC xenograft growth *in vivo*

To determine the effects of liver EC-secreted factors and HER3 inhibition on PDAC tumor growth *in vivo*, we used a proof-of-principle xenograft model with BxPC-3 cells (HER3 +ve). First, we subcutaneously (subQ) implanted BxPC-3 cells in Matrigel into athymic nude mice. Once the tumor burden was confirmed, (10 days after implantation), mice were randomized into four groups with equal average tumor sizes and then treated with CM or/and seribantumab. To replicate the paracrine effect of liver EC-secreted factors on PDAC tumors, CM from human primary liver ECs (EC-9) were injected subQ into the spaces between the tumors and skin tissues once a week throughout the experiment, with BxPC-3 CM as a control. The HER3 antibody seribantumab was delivered by intraperitoneal injection twice a week at 20 mg/kg, a clinically relevant dose used in prior studies (30,41,42). We found that compared to PDAC CM, liver EC CM resulted in an increase of over 2-fold in tumor sizes. Moreover, seribantumab nearly completely stopped tumor growth, resulting in a “stable disease response” in tumor-bearing mice (Fig. 6A, B). Interestingly, seribantumab blocked tumor growth even without EC CM treatment. This finding supports our *in vitro* data that seribantumab decreased cell growth in cells with PDAC CM, as PDAC cells also secrete NRGs for activating HER3 in an autocrine fashion (36). Of note, we also found that EC CM promoted tumor growth in xenografts derived from MIA-PaCa2 cells (HER3 -ve, Suppl. Fig. 4), confirming that liver ECs activate HER3-independent pathway(s) for promoting tumor growth in PDAC cells without HER3 expression.

**Fig. 6.**
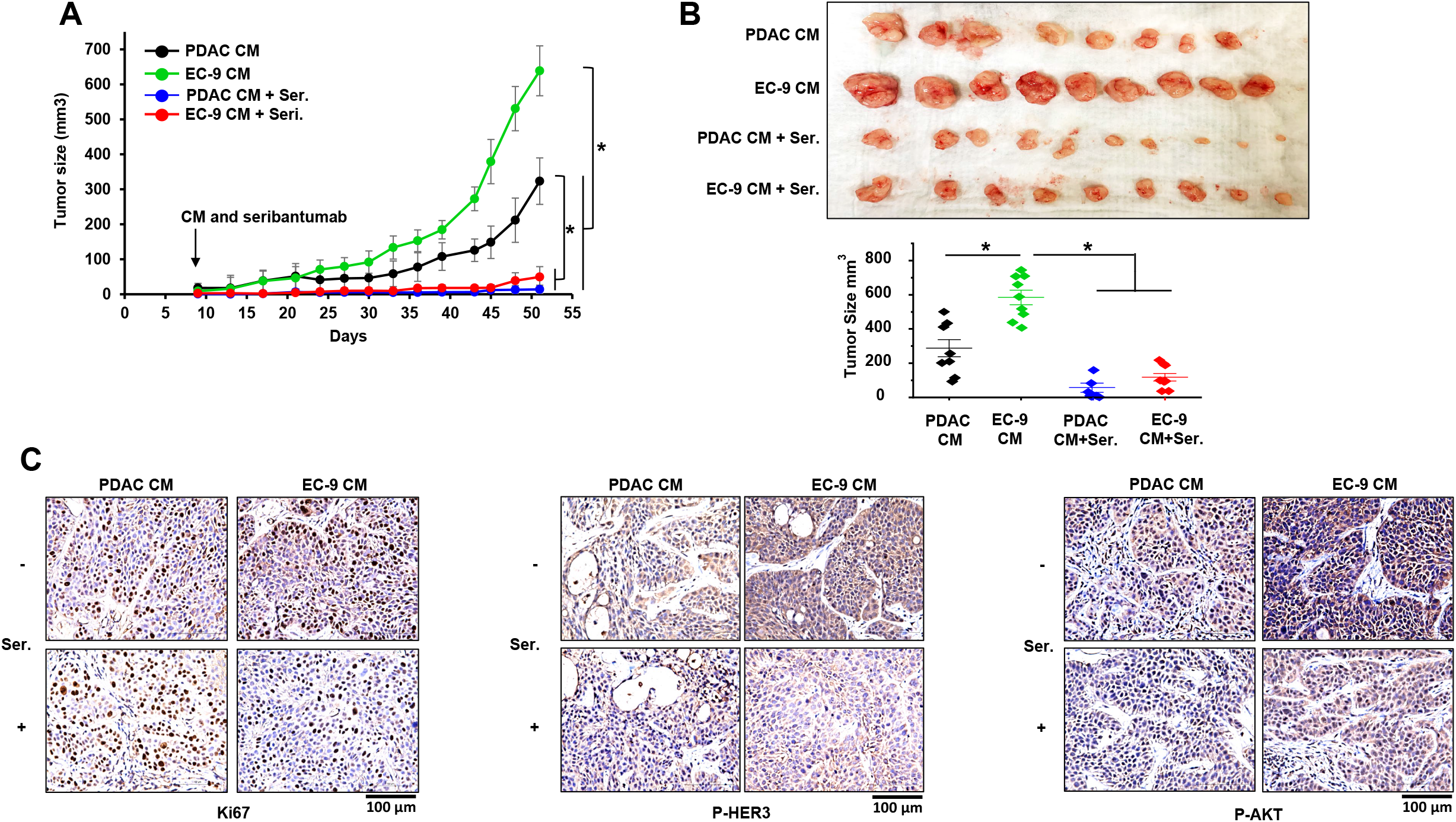
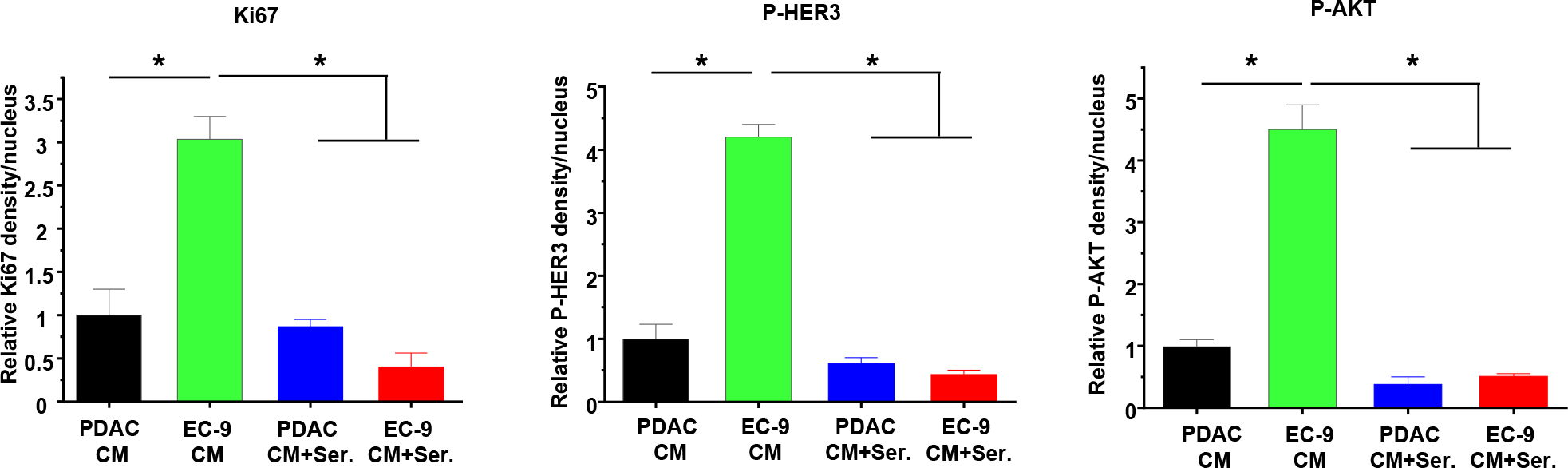
HER3 antibody seribantumab blocked EC-induced PDAC xenograft growth *in vivo*. BxPC-3 cells (HER3 +ve) were implanted subQ. Once tumor sizes were confirmed by a caliper on Day 9, mice were randomized and then treated with CM from BxPC-3 cells (PDAC CM) or liver ECs (EC-9 CM), and with control IgG or seribantumab. **(A)** Tumor size measurements over time show that EC CM increased tumor growth, and that seribantumab blocked EC CM-induced tumor growth. Mean -/+ SD, *P<0.01 one-way ANOVA test between groups after Day 39. **(B)** Image of harvested BxPC-3 xenografts with quantification of tumor sizes by a caliper. *P<0.01 one-way ANOVA test. **(C)** IHC staining of xenograft sections with Ki-67, phospho-HER3 (P-HER3), and phospho-AKT (P-AKT), and were counter stained with hematoxylin. Scale bars represent 100 µm. **(D)** Quantifications of the stained sections (n=3 tumors/group) show EC CM increased levels of Ki-67, P-HER3 and P-AKT in BxPC3 xenograft, all of which were significantly decreased by seribantumab. Mean -/+ SD, *P<0.001 one-way ANOVA test.

To assess the effects of EC CM and seribantumab on HER3-AKT and PDAC cell growth, BxPC-3 xenografts were harvested at the end of the study and processed for IHC staining and quantification of phosphorylation of HER3 and AKT (P-HER3 and P-AKT), and the proliferation marker Ki-67 (Fig. 6C, D). Compared to PDAC CM, EC CM treatment significantly increased the levels of P-HER3 and P-AKT, and the number of Ki-67 positive cells in tumor tissues, and seribantumab significantly decreased the levels of these targets. In agreement with the tumor growth data above, we found that PDAC CM-treated tumors had detectable levels of P-HER3 and P-AKT, both of which were significantly decreased by seribantumab. Taken together, these findings suggest that EC CM activated HER3-AKT in PDAC xenografts and promoted xenograft growth *in vivo*. More importantly, we demonstrated that the HER3 antibody seribantumab significantly blocked tumor growth in HER3 +ve PDAC xenografts, resulting in a nearly stable disease response over the course of 51 days. Such dramatic anti-cancer effect from a drug is rarely seen in a disease that leads to an extremely low survival rate, even in subQ xenograft models.

## Discussion

The regulations of cancer cell functions by stromal cells, mostly fibroblasts and immune cells, in the microenvironment of metastatic tumors have been assessed extensively by others (43,44). ECs are also known to play a role in solid tumor development and metastasis, with much attention on their roles in angiogenesis for establishing blood vessels in and around the tumors. Unlike precedent EC studies focused on angiogenesis and vascular remodeling, studies from our group pioneered the discovery that liver ECs secrete soluble factors to promote CRC tumorigenesis in a paracrine fashion (18,19,32). In this study, we used primary human liver ECs and determined that ECs secreted factors to promote PDAC cell growth in a paracrine fashion. Interestingly, we found that PDAC cells have differential expression levels of HER3, and that EC-secreted NRGs activated HER3 and induced growth in PDAC cells with detectable HER3 expression (HER3 +ve). Moreover, we demonstrated that the HER3-specific antibody seribantumab effectively blocked liver EC-induced HER3 +ve PDAC cell and tumor growth. In contrast, in HER3 -ve PDAC cells, liver ECs promoted PDAC cell growth in a HER3-independent manner, and HER3 inhibition had no impact on EC-induced growth in HER3 -ve cells.

In agreement with our findings, previous studies in PDAC demonstrated that HER3 overexpression and the canonical NRG-induced HER3 activation promote PDAC cell proliferation. When overexpressed in PDAC cells, HER3 interacts with other HER receptors (such as EGFR and HER2) to activate downstream survival pathways without additional ligand stimulation (45,46). Coupled with that, PDAC cells express and secrete NRGs (36), which further activate HER3 and augment the endogenous pro-tumor role of HER3 in an autocrine fashion. As a result, blocking HER3 via gene knockdown or inhibitors decreased PDAC cell growth (37,46). However, most of the preclinical studies focused on the endogenous HER3 signaling in cancer cells without considering the exogenous input from surrounding stromal cells. As a result, blocking endogenous HER3 in PDAC cells by gene knockdown or inhibitors led to significant, but modest, attenuation of PDAC cell growth in preclinical studies (37,45,46). In contrast, in the present study we determined that the surrounding liver EC microenvironment that occurs in PDAC liver metastases secretes high levels of exogenous NRGs to activate the pro-survival HER3 signaling in PDAC cells in a paracrine fashion. More importantly, we demonstrated that in the presence of the clinically relevant liver EC stimulation, blocking HER3 by seribantumab attenuated PDAC cell and tumor growth. Meanwhile, we detected basal levels of HER3-AKT activation even without liver EC stimulation, and found that seribantumab decreased PDAC cell growth. These findings validate that the endogenous NRG-HER3 autocrine signaling occurs in PDAC cells. However, as EC-secreted NRGs further activated HER3-AKT at much higher levels and led to 2-3-fold increases in growth in PDAC cells and xenografts, our findings suggest that the surrounding liver EC microenvironment has the capacity to further activate HER3 and promote growth in PDAC liver metastases. As a previous study reported cancer-associated fibroblasts also secrete NRGs for activating HER3 in PDAC (47), findings from us and others highlight an important role of exogenous NRGs from adjacent non-neoplastic stromal cells for activating the pro-tumor HER3 signaling axis in PDAC cells. Furthermore, the significant anti-tumor effects demonstrated by seribantumab strengthened the notion that HER3-targeted therapies can be effective for treating patients with PDAC liver metastases.

HER3-targeted therapies with inhibitors and antibodies have been assessed in different types of cancer, many of which blocked HER3 and tumor growth in preclinical studies (21,48). Seribantumab, the HER3-specific antibody used in the present study, blocked NRG-HER3 binding and activation and demonstrated strong HER3 inhibition in preclinical studies (18,27,30,31), including a recent study from us in colorectal cancer (17). On the other hand, clinical studies have shown that HER3 inhibitors and antibodies assessed failed to improve patient outcomes in different types of cancer (28,29,49-51). However, most of those clinical studies bear a major flaw: the expression of HER3 in the tumor was not assessed. Data from us and others show that not all tumors express HER3 and only HER3 +ve cancer cells and tumors respond to HER3-targeted therapies. These findings suggest that results from those “failed trials” remain controversial and need further evaluation. On the other hand, recent clinical trials have revealed that HER3-targeted therapies were exceptionally effective in a subset of patients in lung (28), ovarian (49), and breast cancer (29) with *NRG1* gene fusion mutations, which lead to highly activated HER3 in cancer cells. As a result, a clinical trial was initiated to use seribantumab for treating patients with all solid tumors that harbor *NRG1* fusion mutations (CRESTONE NCT04383210). This ongoing trial recently reported a 75% response rate for seribantumab in patients with lung cancer (52), suggesting that HER3-targeted therapies can be effective in the patients with tumors that have active HER3 signaling. In PDAC, HER3-targeted therapies have not been assessed to date. A “pan-HER inhibitor” afatinib caused a 3-month partial response in patients with PDAC with *NRG1* fusion mutations (53-55). Another clinical trial attempted to assess the effects of seribantumab for treating patients with *NRG1* fusions specifically in PDAC (NCT04790695). However, as *NRG1* fusion mutations only occur in less than 0.5% of PDAC (56), this trial only enrolled one patient before termination and had no conclusive outcome (57). Although the effects of HER3-targeted therapies on patients with wild-type *NRG1* PDAC have not been assessed, none of the HER3 +ve PDAC cells used in the present study harbor *NRG1* fusion mutations (determined by gene fusion analysis using CCLE and TCGA databases). As seribantumab demonstrated strong anti-tumor effects in HER3 +ve PDAC cells and led to a “stable response” in BxPC-3 xenografts, our data provide a rationale to assess the effects of seribantumab in patients with PDAC that have wild-type *NRG1*.

Clinical studies reported that HER3 expression correlated with PDAC development and progression (58,59), and is associated with poor overall survival (22,24,60). Meanwhile, data from those studies also indicate that over 50% primary PDAC and nearly 30% metastatic PDAC do not have detectable levels of HER3, suggesting the development of these tumors is likely HER3-independent. Moreover, data from us and prior studies collectively show that there are two distinct groups of PDAC cell lines that either have HER3 expression (HER3 +ve, such as BxPC-3, Capan-2, and CFPAC1) or have low or no detectable HER3 expression (HER3 -ve, such as PANC-1, MIA-PaCa2, and Capan-1) (37,45,61). These data suggest that when HER3 is not expressed, PDAC development is likely relying on alternative survival signaling pathways. Indeed, other pro-survival RTKs, such as IGFR and c-MET, are highly expressed in PDAC cells independent of HER3 expression (23,37). Activation of these pathways promotes PDAC cell proliferation and migration. However, a limited effort to compare HER3 +ve and HER3 -ve PDAC cells responses to different RTK inhibitors did not identify a specific RTK that drives PDAC cell survival in HER3 -ve cells (37). In our case, liver ECs did not activate IGFR or c-MET but did activate insulin receptors, as determined by the RTK array. However, in subsequent validations, we found that insulin receptors were not consistently activated by liver EC CM and blocking insulin receptors had little impact on PDAC cell growth. To date, we have not identified the key signaling pathway(s) that mediate HER3 -ve PDAC development, either without or with liver EC paracrine effects. Our laboratory is currently conducting multi-omics analyses to compare the gene and protein expression profiles between HER3 +ve and HER3 -ve cells without or with liver EC stimulations. These studies will help us to better understand the regulation of the key survival pathway(s) in HER3 -ve PDAC cells but are beyond the scope of this study.

Taken together, the results from this study determined an important role of the liver EC microenvironment in promoting PDAC cell proliferation, and identified that when expressed, HER3 is a key mediator of the EC-induced PDAC cell growth. Furthermore, we showed that the HER3 antibody seribantumab significantly blocked EC-induced cell proliferation and xenograft growth in HER3 +ve PDAC cells. Overall, our findings suggest a potential therapeutic strategy of using HER3 antibodies/inhibitors for treating patients with PDAC liver metastases, and potentially with different stages of PDAC.

## Acknowledgments

This project was supported by the NIH grant CA225756 (R.W.), Case GI SPORE Career Award 5P50CA150964-08, NIH institutional grants P30CA043703 (CWRU CCSG).

| Funder Name | Grant Number | Authors supported |
| --- | --- | --- |
| NIH/NCI | R00CA225756 | R. Wang, M. Wright |
| NIH/NCI | 5P50CA150964-08 | M. Wright |
| NIH | P30CA043703 | Cancer Center Support Grant, Case Comprehensive Cancer Center |

