## supplemental figures for "Liver Endothelium Microenvironment Promotes HER3-mediated Cell Growth in Pancreatic ductal adenocarcinoma"

### *Supplementary Figure 1*

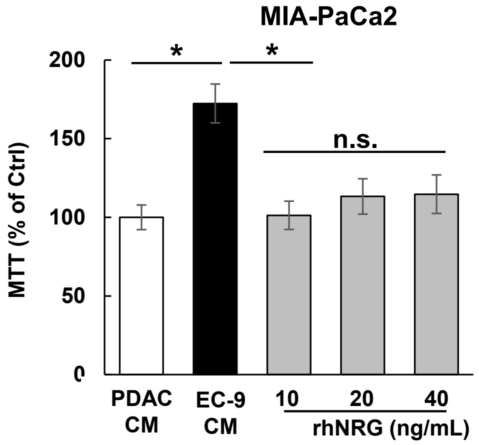


**Supplementary Figure 1. Recombinant human neuregulins (rhNRG) did not increase cell viability in MIA-PaCa2 cells**. MIA-PaCa2 cells (HER3 –ve) were incubated in control PDAC CM, EC-9 CM, or recombinant human NRG (rhNRG) in PDAC CM. The MTT assay showed that different doses of rhNRG had no effect on cell viability, whereas EC CM increased cell viability. Mean +/- SEM of at least three experiments, *p<0.01 t-test compared to control groups with PDAC CM.

### *Supplementary Figure 2*

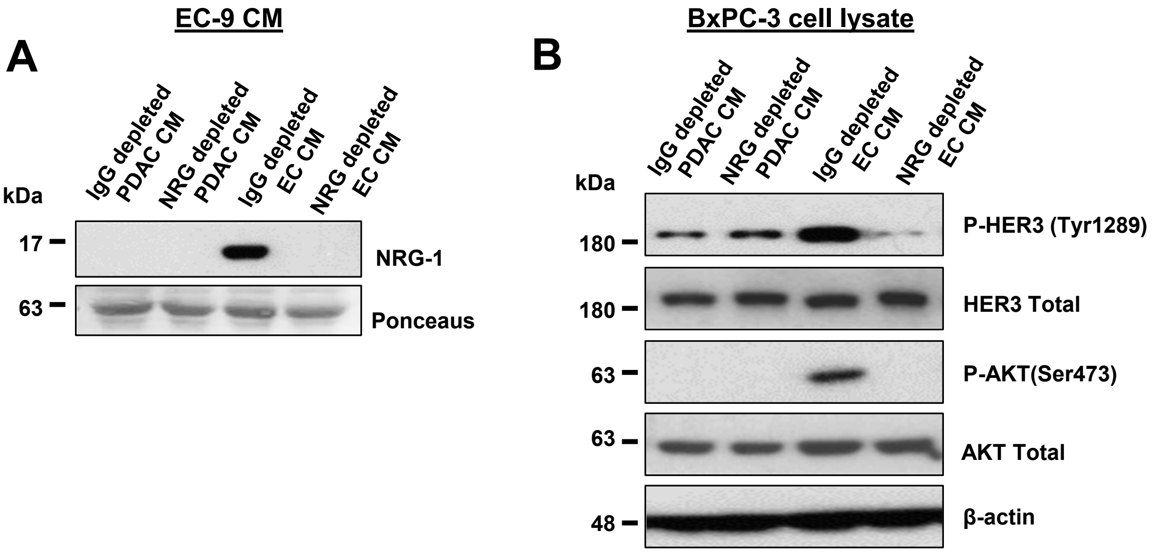


**Supplementary Figure 2. Depleting NRGs from EC CM attenuated EC-induced HER3-AKT activation in PDAC cells.** **(A)** Western blotting showed NRGs were immuno-depleted from EC CM by an NRG-specific antibody. **(B)** Western blotting showed that NRG-depleted EC CM did not activate HER3-AKT in BxPC-3 cells. Total levels of HER3, AKT, and β-actin were used as loading controls. Data represent the results of at least three independent experiments.

#### *Supplementary Figure 3*

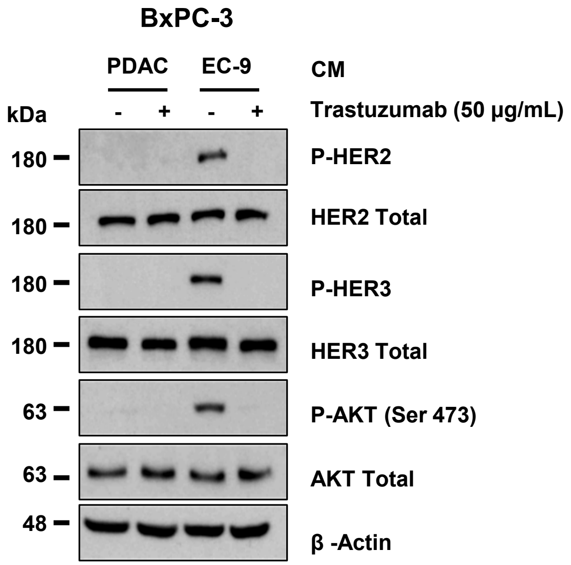


**Supplementary Figure 3. HER2 antibody trastuzumab blocked EC-induced HER3-AKT activation in PDAC cells.** BxPC-3 cells (HER3 +ve) were incubated in control PDAC CM or EC-9 CM and the HER2 antibody trastuzumab (50 µg/mL). The Western blotting showed that trastuzumab blocked liver EC CM-induced HER2, HER3 and AKT phosphorylation. Total levels of HER3, AKT, and β-actin were used as loading controls. Data represent the results of at least three independent experiments.

#### *Supplementary Figure 4*

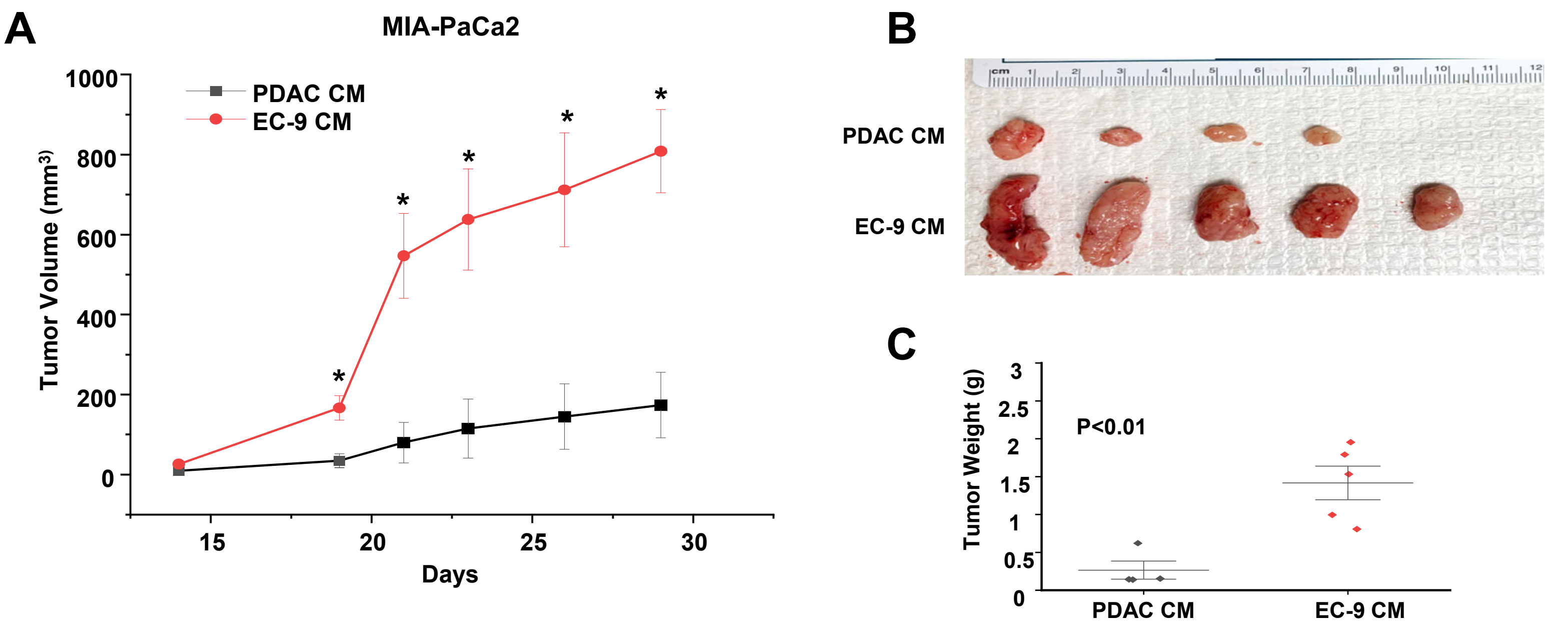


**Fig. 6 EC CM induced PANC-1 xenograft growth *in vivo*.**

MIA-PaCa2 cells (HER3 -ve) were implanted subQ. Once tumor sizes were confirmed by a caliper on Day 10, mice were randomized and then treated with CM from MIA-PaCa2 cells (PDAC CM) or EC-9 CM. **(A)** Tumor size measurements over time showed that EC CM increased tumor growth. Mean -/+ SD, *P<0.01 one-way ANOVA test. **(B)** Image of harvested xenografts. **(C)** Quantification of tumor sizes by a caliper. *P<0.01 one-way ANOVA test.
